# Trade-offs between averages and intra-individual variation within vegetative, phenological, and floral traits

**DOI:** 10.1101/2024.08.30.610450

**Authors:** Charlotte Møller, Martí March-Salas, Mar Sobral, Judith Haase, Pieter De Frenne, JF Scheepens

## Abstract

- Intra-individual trait variation in plants represents an often ignored but important dimension of phenotypic variation that contributes to functional diversity and the dynamics of ecological communities. It can be expressed differently across plant traits, but the induction of intra-individual variation in different trait types under environmental stresses has not yet been explored.
- We used the clonal forest herb, *Galium odoratum*, to investigate intra-individual variation within vegetative, phenological, and floral traits, and trade-offs between trait average and variation under a full-factorial experimental design using drought and shading treatments.
- Intra-individual variation (expressed as CV) differed in magnitude between trait types, with vegetative and floral traits showing the highest and lowest CV, respectively. CV occurring in the different traits responded to the drought and shade treatments. Trade-offs between CV and trait averages appeared across most of traits under the different treatment combinations, whereas trade-offs were less pronounced under control conditions.
- Drought and shading in forest environments induce trade-offs between intra-individual variation and the average trait expression, indicating the relevance of intra-individual variation for functional adaptations of forest plants to climatic changes. Our findings suggest that plastic responses in intra-individual variation may be an important component for mechanistic adjustments of plants to environmental stresses.

## INTRODUCTION

Trait variation occurring within individuals can be referred to as intra-individual variation and is generated by phenotypic plasticity at the organ level in modular organisms – such as plants (Herrera, 2009; Sobral, 2022). Despite its magnitude and ubiquity, intra-individual variation has been largely overlooked, although recent research has shown the relevance of intra-individual trait variation to population dynamics, functional diversity, and ecosystem functioning (Herrera et al., 2022; Sánchez-Bermejo et al., 2024, 2023; Sobral, 2023). However, the mechanisms by which intra-individual variation in different trait types responds to environmental variation, and the functional role of these responses under climate change, remain unknown.

It has been shown that intra-individual variation of vegetative and reproductive traits can be induced by environmental variation (March-Salas et al. 2021; Møller et al., 2023). Furthermore, increased intra-individual variation in floral (Paglia et al., 2023) and phenological traits (Gómez et al., 2022), such as flowering start and longevity, allow plants to enhance their fitness as well as shifting their pollination niche under stressful conditions. Yet, the extent to which different traits (e.g., vegetative growth, phenological responses, reproductive traits, leaf traits and root traits) vary intra-individually and how variation is environmentally induced is not yet well understood.

Within individual plants, variation in vegetative traits, such as leaf size and plant height, has been found to be highly plastic, and affected by environmental cues (Møller et al., 2023). Generally, flowers have been found to be less variable compared to vegetative traits (Herrera, 2009), possibly because floral traits are under stronger selection (e.g., by pollinators) compared to leaf traits (Paglia et al., 2023). On the contrary, one could speculate that leaf traits are more plastic, as they are under weaker selection than floral traits, in response to abiotic selective forces (Brunet et al., 2021; Xiang et al., 2021), such as light and water availability or their combination (Lambrecht et al., 2017; Navas and Garnier, 2002). Moreover, phenological traits, such as flowering start and longevity, rely heavily on micro-environmental cues from their surroundings, being especially sensitive to rising temperatures (Yan et al., 2021; Horbach et al., 2023). Variation in flowering phenology might be beneficial for the individual, as it may allow for a wider range of pollinators or continuously align the onset of flowering with a warming climate, but it may also have detrimental effects, for instance if the activity window of a primary pollinator is missed or if the flowering period includes climatically favourable periods, causing limited capacity for seed production or lack of opportunity to reproduce during the growing season, respectively (Elzinga et al., 2007). Thus, different trait types within individual plants can vary differently in their intra-individual variation under environmental stresses, possibly due to their evolutionary history and plasticity, which may trigger contrasting evolutionary dynamics in plant populations.

High levels of intra-individual variation may be seen as a form of a bet-hedging strategy (Martín et al., 2019), where plants become more efficient at dealing with environmental variability by producing a range of phenotypes, for example within a single clonal individual (Møller et al., 2023). This strategy increases the ability to deal with unexpected environmental stresses, but it may come at the cost of not fully optimizing the average trait response for specific environmental conditions (Haaland et al., 2019). On the contrary, low intra-individual variation will most likely not allow for traits to accommodate for environmental changes, and the traits will more likely be canalized and become fixed (Valladares et al., 2002; Van Buskirk and Steiner, 2009). A trade-off between intra-individual variation and trait averages may reflect a balance between the benefits of maintaining high levels of variability within individuals and the advantages of having optimal (and, depending on the trait, but in many cases high) trait averages within and across individuals (Herrera, 2009; Wetzel et al., 2016).

Trade-offs refer to the variation in one trait to maintain the performance or a specific function of another plant trait when an environmental resource is limited (Stearns, 1998). It was recently shown that intra-individual variation was sometimes associated with the mean response of plant traits in certain environments (Møller et al., 2023). It was also shown that plants can incur different trade-offs between intra-individual variation and trait averages under various environments (Møller et al., 2023). However, it remains hard to interpret the functional role of such observations in plants. For instance, specific environmental cues can change intra-individual variation in flower size without altering the average flower size within an individual plant. Here, the degree of intra-individual variation impacts the functionality of this trait (Sobral, 2023). Higher intra-individual variation in flower size may favour variation in pollinators. This can subsequently cause different fruit sizes, variable fruit sugar contents, and even diversity in time till fruit maturation, which may finally determine the identity and size of the disperser, and later the germination timing and success (Sobral, 2023). Therefore, even population dynamics and persistence could be linked to the plant’s strategy to maintain the mean trait response even when intra-individual variation is altered by changing environmental conditions.

Here, we aim to further our understanding on the extent and relevance of intra-individual variation in different trait types, and on trade-offs between trait averages and intra-individual variation under different environmental conditions. More specifically, we compare intra-individual variation of vegetative, phenological, and floral traits using the herbaceous forest-floor plant *Galium odoratum* (Rubiaceae). Forest understory herbs are increasingly subject to climate-change driven drought (Dai et al., 2018) and altered light regimes due to changed tree leaf-out phenology (Heberling et al., 2019). In this context, we test how experimental drought and the timing of shading, as key global change drivers in European forests (De Frenne, 2024), induce shifts in intra-individual variation in various trait types and in the relationship between trait variation and trait averages. Thus, we ask (1) whether the amount of intra-individual variation varies among different vegetative, phenological, and floral traits; (2) whether this variation and the corresponding trait averages are affected by environmental stress in the form of treatments aimed to simulate climatic changes in forest environments; (3) and how trade-offs between intra-individual variation and trait averages vary under environmental stress.

## MATERIAL AND METHODS

### Study species and experimental system

We used *Galium odoratum* (L.) Scop. (Rubiaceae) as our study species. *Galium odoratum* is a perennial forest forb occurring in Europe, reaching heights of 10-30 cm, and flowering in April-June. Reproduction can occur sexually via seeds, but *G. odoratum* also heavily relies on vegetative spread through stolons (Frederiksen and Rasmussen, 2006). The leaves are lancet-shaped, widest at the middle or just above, and mostly appearing eight per whorl at a time. The flowers are white, small, funnel-shaped and occur in cymes (Lim, 2014). The species’ primary habitat is shaded woodlands with moist and rich soils (Frederiksen and Rasmussen, 2006).

In May 2020, we sampled 5 individuals of *G. odoratum* each from a total of 21 populations (100 m × 100 m forest plots; 105 plants in total) distributed across three regions (i.e. Biodiversity Exploratories, www.biodiversity-exploratories.de) across Germany: Schwäbische Alb (South Germany), Hainich-Dün (Central Germany), and Schorfheide-Chorin (North Germany). Within each region, the sampled plots contrasted strongly in terms of forestry and management intensities as well as in terms of landscape structure and environmental conditions (Fischer et al., 2010). The five individuals per population (hereafter referred to as genets) were sampled with a minimum inter-individual distance of 10 m to ensure sampling genetically different individuals (Ziegenhagen et al., 2003). Each genet was vegetatively propagated into four ramets and transplanted in multitrays (51.5 cm wide, 33.5 cm long, each pot 5.0 cm diameter and 5.5 cm deep, 54 pots per tray), filled with potting soil (CL T torffrei, Einheitserde, Sinntal-Altengronau, Germany) and placed in an outdoor common garden for establishment and growth until November 2020. Subsequently, all ramets were transferred into 1.5 L pots with potting soil (Typ T, Struktur 1B, Hawita, Vechta, Germany) and with trays placed underneath. In spring 2021, we relocated all pots to a foil tunnel with open ends to exclude natural precipitation. After the first year, but before we took measurements for this study, 65 out of 105 genets from the 21 populations survived and were used for phenological and floral measurements. Vegetative measurements were taken after the flowering peak, where some additional mortality had occurred, resulting in a total of 425 ramets deriving from 51 genets from 15 populations (see Table S1 and Table S2).

### Experimental shade and drought treatments

Shading and drought treatments were applied in 2021 and 2022 in a full-factorial design, meaning that one of the fours ramets per genet was placed under each of the four different treatment combinations: 1) control, 2) drought, 3) early shading, and 4) combined drought and early shading treatment. To simulate the two-week shaded forest understorey environment and climate-induced mismatch between forest over- and understorey, we applied shading cloth over the foil tunnel in two layers (45% shading for each layer, resulting in 90% total shading). The two shading cloth layers were applied one week apart. The first layer of the early shading treatment was applied ∼2 weeks before the anticipated leaf out of beech and oak trees in the surrounding area (that is, on 12 April in 2021 and 11 April in 2022). Control shading cloth was applied when the natural leaf-out of surrounding trees was observed in the area (Frankfurt am Main, Germany) on 30 April in 2021 and 25 April in 2022 (again the two layers were applied one week apart). The drought treatment was applied to all plants when the first plant started flowering (on 7 May in 2021 and 30 May in 2022). This drought treatment was applied as a single event, in which all watering ceased until 50% of the plants were wilting (which took approximately three weeks). At this point regular watering *ad libitum* resumed for all pots. Control plants regularly received water *ad libitum* during the whole experimental period. Watering was done through irrigation from above.

### Mean and intra-individual trait measurements

Measurements for this study were taken in 2022. The measured vegetative traits included leaf length and leaf width, measured with callipers to the nearest 0.1 mm precision on one randomly chosen leaf on the top leaf whorl on up to five ramets per pot. The measured phenological traits included start of flowering and flower longevity and the measured floral traits included flower size and petal size. Flowering start (defined as when the anthers became visible) was noted down as day 1 when the first flower started flowering, and subsequently checked every Monday, Wednesday, and Friday between 1^st^ of April and 1^st^ of June 2022, for the first 15 flowers per ramet. Two days after flowering start, the flower size (diameter of the flower) and petal size (length of a petal) were measured with callipers to the nearest 0.1 mm precision for the same 15 flowers. Furthermore, the flower longevity, that is the total number of days that a flower was open until the wilting of flower petals, was tracked every Monday, Wednesday, and Friday between 1^st^ of April and 6^th^ of June 2022. We measured a total of 425 leaves originating from a total of 135 ramets (average number of leaves ± standard deviation per pot: 4.9 ± 0.28; see Table S1 for the distribution of the vegetative trait measurements across the treatments), and 1621 flowers originating from the same 135 ramets (average number of flowers ± standard deviation per ramet: 14.38 ± 3.64; see Table S2 for the distribution of the floral trait measurements across the treatments).

As an estimation of intra-individual variation, the coefficient of variation (CV) was calculated as the standard deviation divided by the mean of a specific trait at the intra-ramet level. We only included leaf measurements from the top whorls of *G. odoratum* shoots, as these are closest to the flowers, controlling for effects of plant development and spatio-temporal variation in the microenvironment when comparing floral and leaf traits.

### Data analyses

All statistical analyses were conducted with R version 2023.09.1 (R Core Team, 2023).

First, we tested for the effects of the treatments on the CV for all traits combined using an LMM. The CV calculated within each trait in the complete dataset (CV_ALL)_ was used as the response variable, with the trait average within each trait as covariate, as well as the traits (flowering start, flower longevity, flower size, petal size, leaf length, or leaf width) and treatments (control, drought, early shade, and combined drought and early shade) as fixed factors. As random factors we included genet nested within population nested within region (region/population/genet). The LMM was run using the ‘lmer’ function from the ‘lme4’ package (Bates et al., 2009), and LMM test results were obtained by applying the ‘Anova’ function from the ‘car’ package (Fox et al., 2007). Significant effects of trait identity and the treatments on CV_ALL_ were further investigated with a post-hoc Tukey test using the ‘emmeans’ function from the ‘emmeans’ package (Lenth and Lenth, 2018).

Second, we also tested for the effects of the treatments on each specific trait separately. Here, the trait CV was used as response variable, with trait average, shade and drought treatments and their interaction as fixed factors. Again, genet nested within population, nested within region, were used as random factors. Normality and homoscedasticity of all model residuals were evaluated visually with qq-plots and histograms. When at least one of these assumptions was violated, the response variable was transformed using square-root transformation. Significant effects of treatment interactions on CVs were further investigated with post-hoc Tukey tests using the ‘emmeans’ function from the ‘emmeans’ package (Lenth and Lenth, 2018). The same analyses were repeated with the trait averages as response variables and trait CV as a fixed factor.

Third, we performed a principal component analysis (PCA) using the ‘prcomp’ function to identify dimensions of intra-individual trait variability and of trait averages and to investigate potential trade-offs between CV and trait averages. We analysed the data for each treatment independently: control, drought, early shading, and combined drought and early shading, encompassing 26, 33, 28, and 26 individuals, respectively. Each individual trait CV and trait average was log-transformed and scaled for appropriate comparison. Furthermore, the PCA loadings, representing the trait CVs and trait averages, were extracted for the first two axes for each treatment PCA (Control, Fig. S1; drought, Fig. S2; early shading, Fig. S3; combined, Fig. S4).

## RESULTS

Intra-individual variation, assessed as CV_ALL_ within ramets across all traits, differed significantly among traits (Table 1; Fig. 1). Flower size (mean CV ± Standard Error: 8.8 ± 4.1 SE), petal size (9.2 ± 4.1 SE), and flowering start (11.2 ± 6.3 SE) displayed the lowest CV. Flower longevity (24.1 ± 10.4 SE) had intermediate CV, and leaf width (33.4 ± 14.6 SE) and leaf length (37.9 ± 14.8 SE) had the highest CV out of the measured traits (Fig. 1). Furthermore, CV_ALL_ was significantly negatively related to the corresponding overall trait averages, although the effect was weak (Table 1; Parameter estimate: -0.03). Furthermore, the corresponding trait averages significantly affected CV_ALL_ (Table 1).

**Figure 1.**
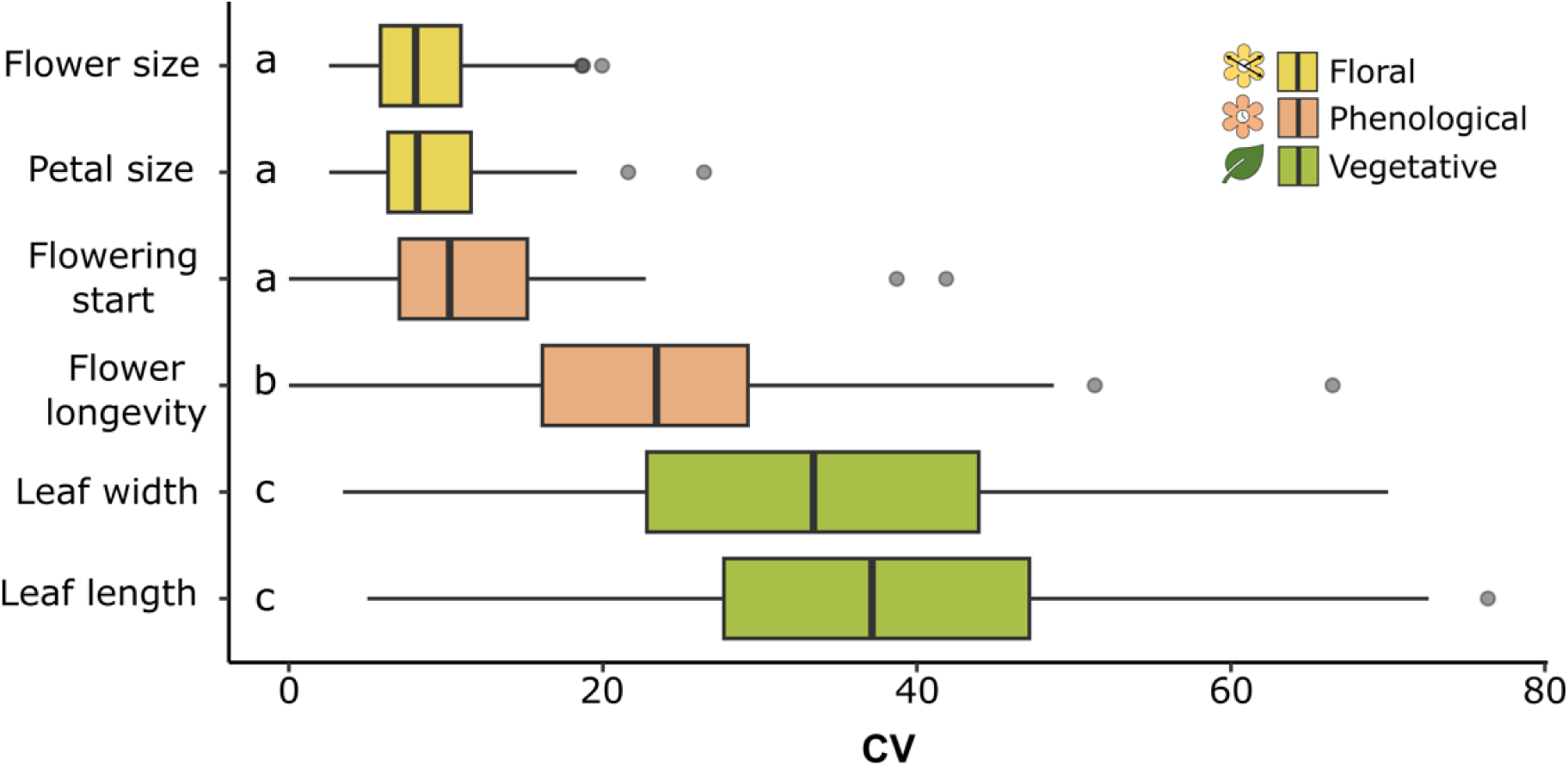
Intra-individual coefficient of variation (CV) of flower size, petal size, flowering start, flower longevity, leaf width, and leaf length at the intra-ramet level derived from the complete dataset (CV_ALL_). The trait types (floral, phenology, and vegetative) are indicated by colour and icon. Significant differences based on a post-hoc Tukey test among the traits are indicated with letters.

**Table 1.**
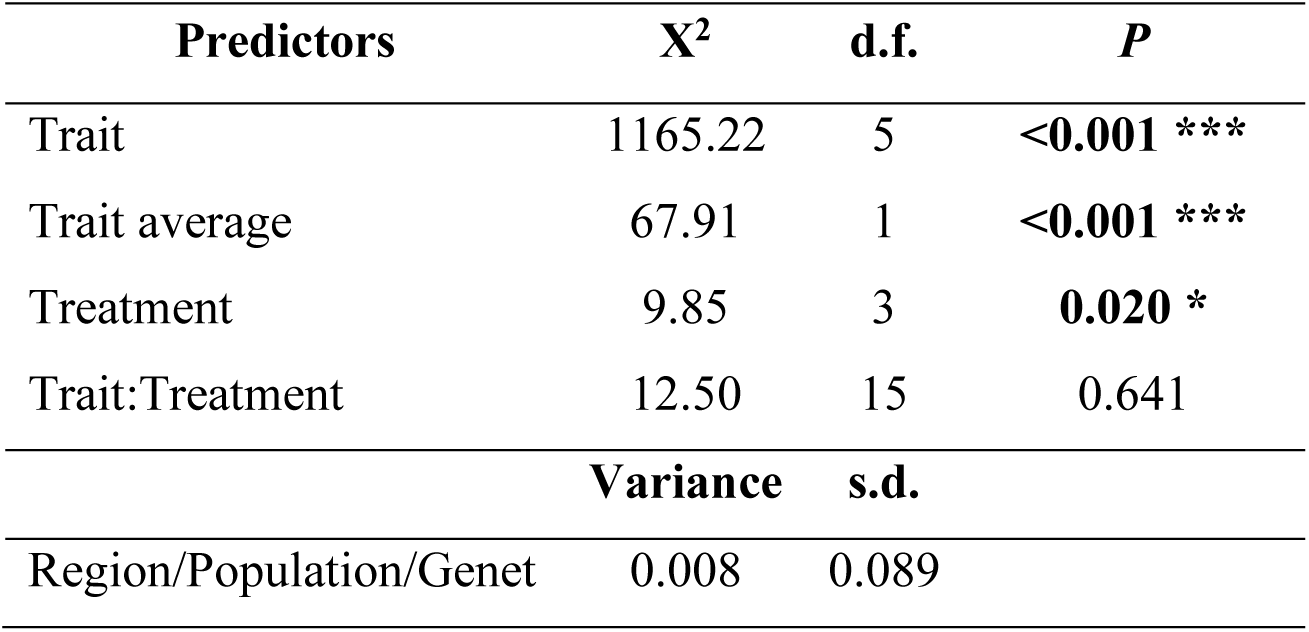
Results of the mixed-effects model of the intra-individual coefficient of variation (CV) calculated within each trait in the complete dataset (CV_ALL_) in *Galium odoratum* at the intra-ramet level as response variable. Trait, average trait value, the experimental treatments and the interaction between trait and treatment were included as fixed factors. Genet nested within population and region was included as random factor. Variance and standard deviation (s.d.) are provided for the random factor. Significance levels are represented by asterisks: * *P* < 0.05; ** *P* < 0.01; *** *P* < 0.001.

We found a significant effect of the treatments on both CV_ALL_ (Table 1), as well as on some trait specific CVs (Table 2). More specifically, the CV of flower and petal size was highest under control conditions (Fig. **2a**, **2b**), lowest under combined treatment conditions and intermediate under drought and early shading. In comparison, averages of floral traits were mostly of equal size across treatments, except for under the early shading treatment, where flower and petal size tended to be slightly higher (Fig. **2a**, **2b**). Furthermore, CV of flowering start tended to increase CV under drought and early shading (Fig. **2c**). In comparison, average flowering start was similar across treatments, except for under the drought treatment, where individuals tended to flower earlier (Fig. **2c**). No trends in CV of flower longevity among treatments could be observed (Fig. **2d**). However, mean flower longevity tended to be slightly higher under the drought treatment compared to control, early shade, and the combined treatment (Fig. **2d**). Drought conditions appeared to reduce the CV of leaf length and leaf width, while the combined treatment increased the CV (Fig. **2e**, **2f**). In comparison, vegetative trait averages were also found to decrease under all treatments compared to the control (Fig. **2e**, **2f**).

**Figure 2.**
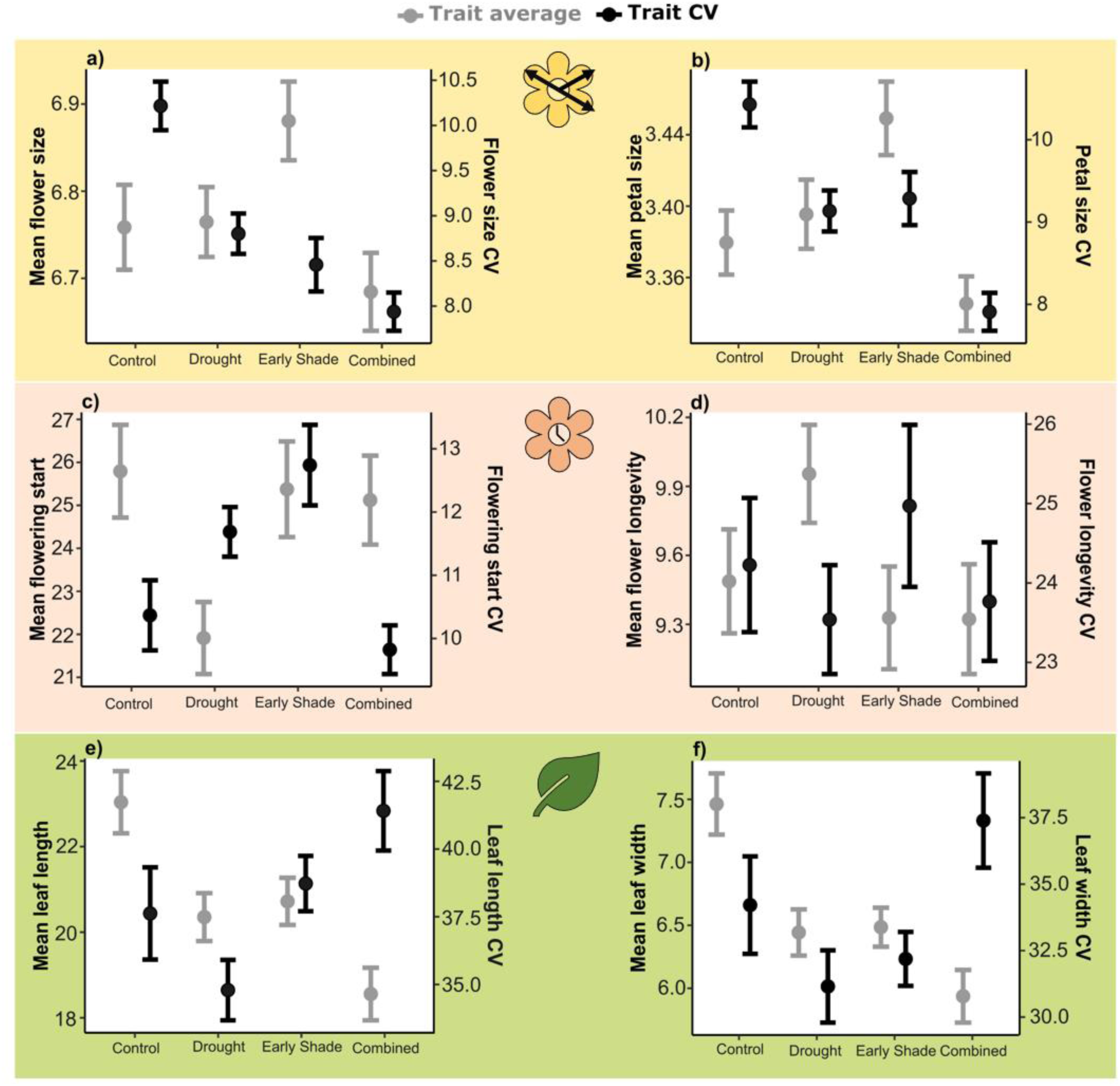
Treatment effects on the predicted intra-individual variation (CV) and predicted means of **a)** flower size, **b)** petal size, **c)** flowering start, **d)** flower longevity, **e)** leaf length, and **f)** leaf width at the intra-ramet level in *Galium odoratum*. The trait types (floral, phenology, and vegetative) are indicated by colour and icon.

**Table 2.** Effects of the applied treatments on trait-specific variation at the intra-ramet level. Coefficient of variation (CV) in *Galium odoratum* at the intra-ramet level calculated for each trait was used as response variables. Trait average as well as the experimental treatments and their interaction were included as fixed factors and covariates. Genet nested within population and region was included as random factor. Significance levels are represented by asterisks: • *P* <0.1; * *P* < 0.05; ** *P* < 0.01; *** *P* < 0.001.

| Response | Predictor | Chisq | Df | P |
| --- | --- | --- | --- | --- |
| Flowering start CV | Trait average | 0.378 | 1 | 0.539 |
|  | Shade | 0.178 | 1 | 0.673 |
|  | Drought | 1.249 | 1 | 0.264 |
|  | Shade:Drought | 1.570 | 1 | 0.210 |
| Flower longevity CV | Trait average | 18.009 | 1 | **<0.001 \*\*\*** |
|  | Shade | 0.038 | 1 | 0.845 |
|  | Drought | 0.354 | 1 | 0.552 |
|  | Shade:Drought | 0.184 | 1 | 0.668 |
| Flower size CV | Trait average | 9.102 | 1 | **0.003 \*\*** |
|  | Shade | 3.115 | 1 | **0.078 •** |
|  | Drought | 2.615 | 1 | 0.106 |
|  | Shade:Drought | 0.204 | 1 | 0.652 |
| Petal size CV | Trait average | 7.781 | 1 | **0.005 \*\*** |
|  | Shade | 3.755 | 1 | **0.053 •** |
|  | Drought | 4.341 | 1 | **0.040 \*** |
|  | Shade:Drought | 0.056 | 1 | 0.807 |
| Leaf length CV | Trait average | 11.433 | 1 | **<0.001 \*\*\*** |
|  | Shade | 0.017 | 1 | 0.896 |
|  | Drought | 1.450 | 1 | 0.229 |
|  | Shade:Drought | 0.186 | 1 | 0.666 |
| Leaf width CV | Trait average | 14.633 | 1 | **<0.001 \*\*\*** |
|  | Shade | 0.098 | 1 | 0.754 |
|  | Drought | 1.193 | 1 | 0.275 |
|  | Shade:Drought | 2.353 | 1 | 0.125 |

The PCAs show that under the control treatment (Fig. **3a**; Fig. S1), trait CVs and trait averages are defined almost equally by PC1 (31.4%) and PC2 (25.4%), and a negative relationship (opposing arrows) between trait CVs and trait averages can be observed for flower longevity, leaf length, and leaf width (Fig. **3a**; Table 3). Under drought (Fig. **3b**), the averages of the floral traits (flower size and petal size) define the first axis (25.2%), whereas the CVs and averages of vegetative traits (leaf width and length) define the second axis (22.4%) as well as the fourth axis (10.8), and lastly, flower longevity CV defines the third axis (14.4%) (Fig. S2). More specifically, in the drought treatment we observe a negative relationship between CVs and trait averages in all traits except for flowering start, which had no correlation (Fig. **3b**; Table 3). Under early shading (Fig. **3c**), floral trait CVs and averages defined the first axis (38.7%), and phenological trait averages were the most important traits on the second axis (18.3%). No clear relationship can be found on the third axis (14.1%), but vegetative trait averages seem to dominate the fourth axis more extensively (12%) (Fig. S3). Furthermore, we observe a negative relationship between the CVs and averages of floral traits and of flowering start under early shading, respectively (Fig. **3c**; Table 3). Lastly, under combined treatments (Fig. **3d**), vegetative trait CVs defined the first axis (33.5%), while the second axis (23.2%) was dominated by floral trait CVs and averages. The third axis (13.3%) mainly consists of vegetative trait averages, and lastly, the fourth axis (10.1%) was defined by flower size CV as well as flower and petal size trait averages (Fig. S4). Here, we observe negative relationships between CVs and averages along PC1 for both vegetative traits and for flower longevity, and along PC2 for floral traits, whereas flowering start CV and average did again not show any correlation (Fig. **3d**; Table 3).

**Figure 3.**
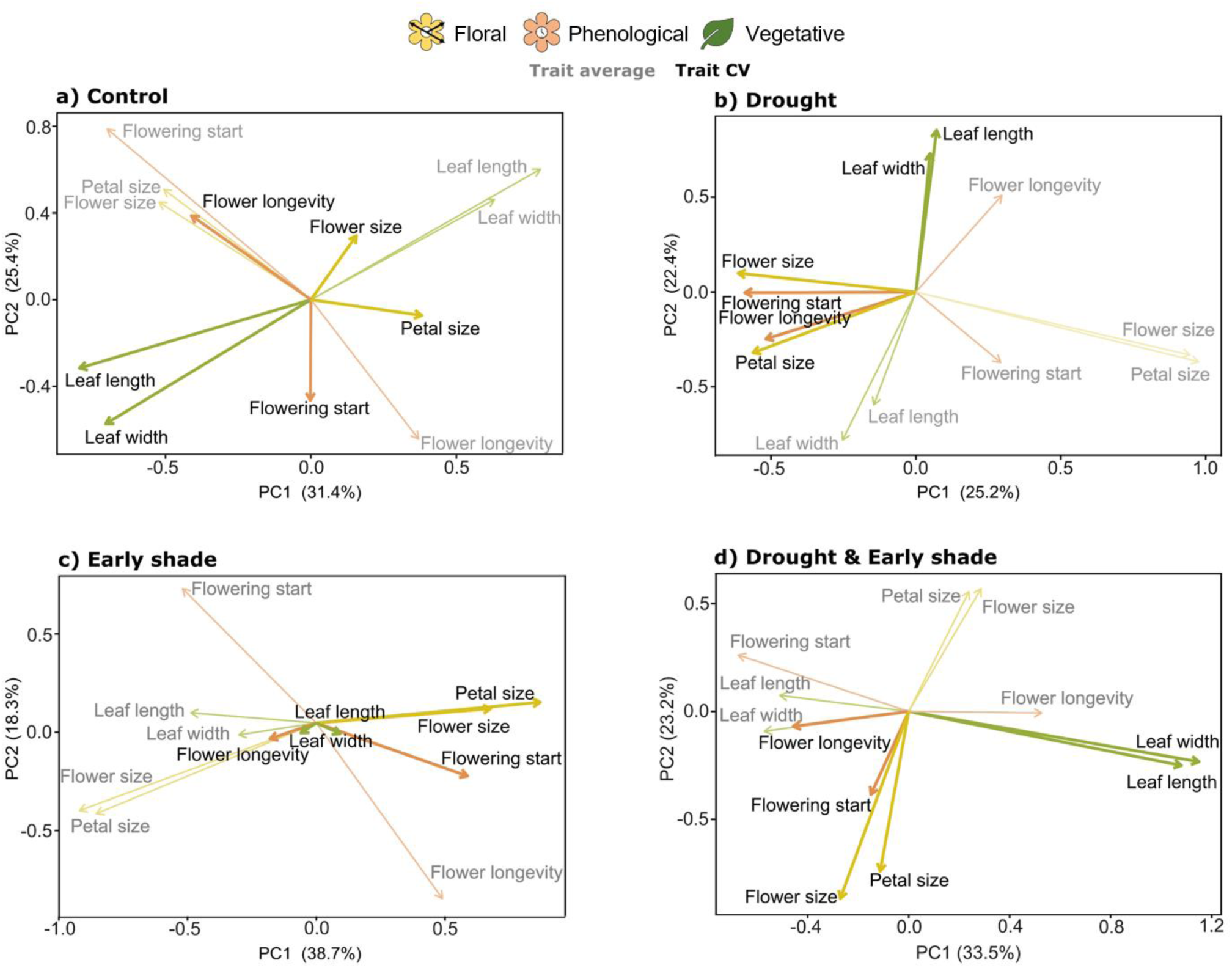
Principal component analysis (PCA) of average trait values (grey text and arrows of lighter colours) and intra-individual variation (CV; black text and arrows of darker colours) for **a)** the control treatment, **b)** drought treatment, **c)** early shading treatment, and **d)** the combined drought and early shading treatment. In all cases, the first two axes (PC1 and PC2) are depicted and the amount of explained percentage is indicated. Solid arrows represent the direction of the studied trait CVs and averages. Trait types (floral, phenology and, vegetative) are indicated by colour (arrows) and icon.

**Table 3.** Overview of trade-off relationships. Positive correlations (+; <60ᵒ), no correlations (×; 60-120ᵒ), and negative correlations (−; >120ᵒ) between trait CV and trait averages based on results shown in Figure 3.

| Trait CV-average trade-off | Control | Drought | Early shade | Drought &<br>Early shade |
| --- | --- | --- | --- | --- |
| Flower size | × | – | – | – |
| Petal size | × | – | – | – |
| Flowering start | × | × | – | × |
| Flower longevity | – | – | × | – |
| Leaf length | – | – | × | – |
| Leaf width | – | – | × | – |

## DISCUSSION

Here, we show that intra-individual variation (expressed as coefficient of variation – CV) varies strongly among traits, with the lowest amounts of variation found in floral traits, intermediate amounts in phenological traits, and the highest amounts in vegetative traits. Trait CV was occasionally found to be influenced by experimental drought, early shading, or their interaction, with the direction of the response depending on the trait. Furthermore, we show that trade-offs between trait CVs and averages are impacted by the treatments, with some trade-offs disappearing or appearing under particular treatments, and with all observed correlations between trait CV and averages being negative. These findings suggest that plastic responses in intra-individual variation may be an important driver for adjusting plant responses to environmental change.

### Differences in intra-individual variation among plant traits

The measured traits differed in the extent of exhibited intra-individual variation (CV). We found the highest amount of intra-individual variation in the vegetative traits leaf length and leaf width, concurrent with previous findings (Herrera, 2009). Larger variation in leaf traits may allow individuals to thrive under variable light and drought conditions, simultaneously increasing light capture ability while decreasing potential evapotranspiration (Møller et al., 2023). Flower traits such as petal and flower size were found to have low levels of intra-ramet CV. Flower size CV is a trait that has been observed to be under selection exerted by pollinators (Herrera, 2009; Parachnowitsch and Kessler, 2010; Caruso et al., 2019), offering a potential explanation for the little variation within flowers. Small variation in floral traits has previously been shown to have important evolutionary consequences for pollinator interactions, albeit only on the inter- and intra-specific level (Lambrecht and Dawson, 2007). Oppositely, increased flower size variation within an individual could accommodate for an increased number of pollinator species (Gómez et al., 2020). Low amounts of intra-individual variation were also observed for flowering start, whereas flower longevity was more variable. Timing and response to environmental cues in phenological traits such as flowering start are crucial to retain fitness (Elzinga et al., 2007; Pau et al., 2011). Hence, the low intra-individual variation for flowering start observed in our common garden might be the result of strong selection for simultaneous flowering when pollinators are around and potential risks (such as late frosts) are low. In contrast, the intermediate amount of intra-individual variation in flower longevity suggests that, while flowering start is under strong genetic constraints, individuals can vary flower longevity per individual flower, which may be triggered by the environment. For instance, flowers may finish anthesis after successful pollination (Primack, 1985; Trunschke, 2017).

### Treatment effects on trait averages and CV

Both parallel effects but also contrasting direction and magnitude of treatment effects on trait CV and trait averages could be observed. In vegetative traits, the drought treatment led to reduced CV. One explanation could be that genetic and developmental constraints on leaf growth and morphology become apparent under drought stress (Aroca, 2012; Farooq et al., 2009), which may limit the capacity for high CV in leaf traits. A high CV could be observed in tandem with low trait averages under the combined treatment in vegetative traits. As opposed to the drought treatment, the increased CV in vegetative traits under the combined treatment might be a result of diversification of resource allocation. Leaves developing in more sun-exposed conditions may remain relatively small and thick to reduce the risk of water stress (McDowell et al., 2008; Nardini et al., 2014), while leaves developing in more shady conditions may enlarge leaf area to maximize light capture (Weijschedé et al., 2006).

In floral traits, trait CV was highest under control conditions and decreased with progressively stressful conditions. Under benign conditions, resources allocated to floral traits might allow for great variation in flower and petal size, accommodating for a greater range of pollinators, in the same way that greater variety of fruit sizes attracts various dispersers (Sobral, 2023). Under early shading, trait averages of floral traits slightly increase, enhancing the floral display. This may help attract pollinators, which are often scarce in the forest understorey while competition for pollinators among plants is high. However, under the combined treatment, both trait CV and trait averages in floral traits are at their lowest. These differentiated responses to the various treatments, whether passive or active, could indicate that flower and petal size variation and average might not be a priority for the individual under severe resource constraints, although this interpretation needs further evidence.

Regarding phenological traits, we mainly observed low flowering start averages, and thus earlier flowering, in individuals exposed to drought conditions, which could indicate a drought escape attempt (Shavrukov et al., 2017). However, flowering start showed a more enigmatic pattern in its CV, with low variation in the two extremes, the control, and the combined treatment, and with high variation under drought and early shading conditions separately, which we cannot explain. Finally, neither distinct patterns in flower longevity CV nor in its average could be observed, indicating that longevity might be a constrained trait with little to no plasticity available.

Taken together, we show that intra-individual variation responds to simulated future climatic conditions in the majority of our measured traits and is therefore inducible, highlighting its potential importance for rapid adjustment to the environment and for organismal fitness.

### Trade-offs between trait averages and trait CV

Our results indicate that intra-individual variation is an important component that affects plant strategies under environmental stress. Trade-offs between intra-individual variation on and trait averages may reflect genetic or ecological constraints or adaptive strategies and are so far not well understood. In our study, such trade-offs were evident in all traits, with some trade-offs disappearing or appearing under specific treatments.

Under the control treatment, CVs and averages of floral traits and of flowering start were uncorrelated, implying that under benign conditions individuals are unconstrained in their expression of trait variation vs trait average. In contrast, the negative relationships between vegetative trait CVs and averages reflect that vegetative leaf traits of *G. odoratum* are constrained. This constraint may be due to intrinsic light limitations in forests. Here, the plants either expressed a greater prevalence of smaller leaves, which are less costly to produce and maintain in tandem with a high CV, or strictly expressed large trait averages. The trade-off between trait CV and average in flower longevity could indicate that, under benign control conditions, drought conditions and the combined treatment alike, individuals might either favour longer flower longevity over increased CV or increased CV at the expense of flower longevity, which may allow some flowers to escape from adverse conditions.

Under the drought and the combined treatments, the majority of plant traits were subjected to trade-offs between trait CV and average. First, we observed an opposing relationship between floral trait CVs and averages. Low variation in flower and/or petal size can potentially limit the pollinator niche, at the cost of prioritizing an earlier flowering peak. This could either be viewed as an adaptive response and an attempt to increasing their chances of early pollinator interactions to facilitate a drought escape (Kooyers, 2015; Shavrukov et al., 2017), while it is also possible that it is a passive effect of the drought. Interestingly, some trade-offs appear and disappear depending on the treatment, while other trade-offs are constant among some of the treatments. For instance, the trade-off between CV and trait average in leaf length and leaf width did not change under drought and the combined treatment compared to the control conditions but disappeared under early shade. Under early shading conditions, we observe a trade-off between trait CV and trait averages in the floral traits. On one hand, increased floral display may be particularly useful to attract more pollinators, whose activity could be limited in the shaded conditions (Caudill et al., 2017; Supriyadi et al., 2020). While on the other hand, increased variation in flower display might cater towards a broader range of pollinators in already limited availability due to the shaded conditions. Under combined drought and early shading conditions, we observed strong opposing relationships between trait CVs and trait averages for each trait except flowering start. The applied treatments might be opposing forces for leaf adaptive response. Leaves often decrease in size under drought conditions to limit evapotranspiration, while they can increase under shaded conditions to increase light capture. Perhaps, the increased CV serves to cover several important leaf functionalities such as leaf capture and water retention under these limiting conditions (Milla et al., 2008), which cannot be gained simultaneously with a large trait average. And lastly, similarly to the separate drought and early shading treatments, we again observed a strong negative relationship between floral trait CVs and averages under combined treatments, and as previously suggested increased flower display serves to attract more pollinators, while increased variation in flower display can cover a broader range of pollinator species.

Ultimately, the consistently strong negative relationships observed for the majority of the traits under the drought and early shading treatments, and their interaction, indicate that our simulated future climatic conditions in this experiment alter the plants’ responses, implying that both changes in CV as well as trait averages are key components in plant strategies. Based on our findings, we can speculate that while too little intra-individual trait variation could put an individual at a disadvantage when faced with environmental change, too much variation could balk the functions of each trait type and local adaptations (Kawecki and Ebert, 2004) from solidifying. However, despite putting forth various potential explanations, the functional and ecological meaning behind the distinct trade-off patterns remains unknown, calling for further experimental studies.

### Future directions

Though our current experiment has provided valuable insights, we cannot say for certain whether the observed patterns are caused by adaptive responses or passive responses to the applied treatments. Our study lacks a direct measure of fitness, which is crucial for determining the adaptive significance of the observed patterns, and direct measurements of the quantity and diversity of floral visitors will help to understand underlying ecosystem functions. For *G. odoratum*, suitable fitness measures could be in the form of flowers or seeds produced or number of ramets. Including these metrics in future experiments will enable us to assess whether the observed patterns are the adaptive function of intra-individual variation on its own or merely a passive response. Furthermore, expanding and conducting long-term studies will provide a more comprehensive understanding of how the observed patterns develop and persist over time. This can help to determine whether the responses are consistent and stable indicators of adaptive strategies or if they fluctuate with environmental change.

### Conclusions

In sum, the magnitude of intra-individual variation varied greatly among plant traits in our study species. The lowest values were found in floral traits, followed by phenological traits, and the highest values in vegetative traits, which reflects how intra-individually plastic or constrained different traits and trait types may be under ongoing environmental stresses. Furthermore, intra-individual variation occurring in the different traits was found to be environmentally inducible, as it responded to the drought and shade treatments, indicating the value of intra-individual trait variation to deal with current changes in forest and other environments. Furthermore, our study shows that the trait CV originating from distinct plant traits and trait types are dominant dimensions of our trait functionality measurement spectrum alongside trait averages. Lastly, the ecological meaning behind the observed trade-offs remains unknown but could potentially indicate that plant strategies and trait functionalities can vary depending on the environment, as the intra-individual trait response change across environments. Understanding the extent of variation within different traits and trade-offs involving intra-individual variation is crucial for predicting functional strategies that plants might utilize in response to future climatic changes, and sheds light on potential implications for population evolutionary trajectories and ecosystem dynamics.

## ACKNOWLEDGEMENTS

We thank the managers of the three Exploratories, Julia Bass, Max Müller, Anna K. Franke, Miriam Teuscher, Robert Künast, Franca Marian, and all former managers for their work in maintaining the plot and project infrastructure; Christiane Fischer and Victoria Grießmeier for giving support through central office; Andreas Ostrowski for managing the central data base, and Markus Fischer, Eduard Linsenmair, Dominik Hessenmöller, Daniel Prati, Ingo Schöning, Francois Buscot, Ernst-Detlef Schulze, Wolfgang W. Weisser and the late Elisabeth Kalko for their role in setting up the Biodiversity Exploratories project. We thank the administration of Hainich National Park, the UNESCO Biosphere Reserve Swabian Alb and the UNESCO Biosphere Reserve Schorfheide-Chorin as well as all landowners for their excellent collaboration. We thank Oliver Bossdorf for providing facilities during the initial phase of the project in Tübingen, and Robert Anton, Susanne Pietsch, and the Wissenschaftsgarten at Goethe University Frankfurt for their general support. Lastly, we would like to thank Lutz Stübing, Lena Reimann, Pascal Karitter, Silas Büse and Delia Gärtner for their help in constructing the experimental system and assisting in data collection.

## AUTHOR CONTRIBUTIONS

CM, MMS, PDF, and JFS designed the study. CM conducted the field sampling and JH took the measurements in the common garden experiment. CM, MMS, and MS analysed the data. CM wrote the first draft of the manuscript with all co-authors contributing to revisions.

## DATA AVAILABILITY STATEMENT

This work is based on data elaborated by the HerbAdapt project of the Biodiversity Exploratories program (DFG Priority Program 1374). The datasets are available upon request in the Biodiversity Exploratories Information System under the links: https://www.bexis.uni-jena.de/ddm/data/Showdata/31762 and https://www.bexis.uni-jena.de/ddm/data/Showdata/31766

