## Supplementary Information for "Trade-offs between averages and intra-individual variation within vegetative, phenological, and floral traits"

SUPPLEMENTARY MATERIAL

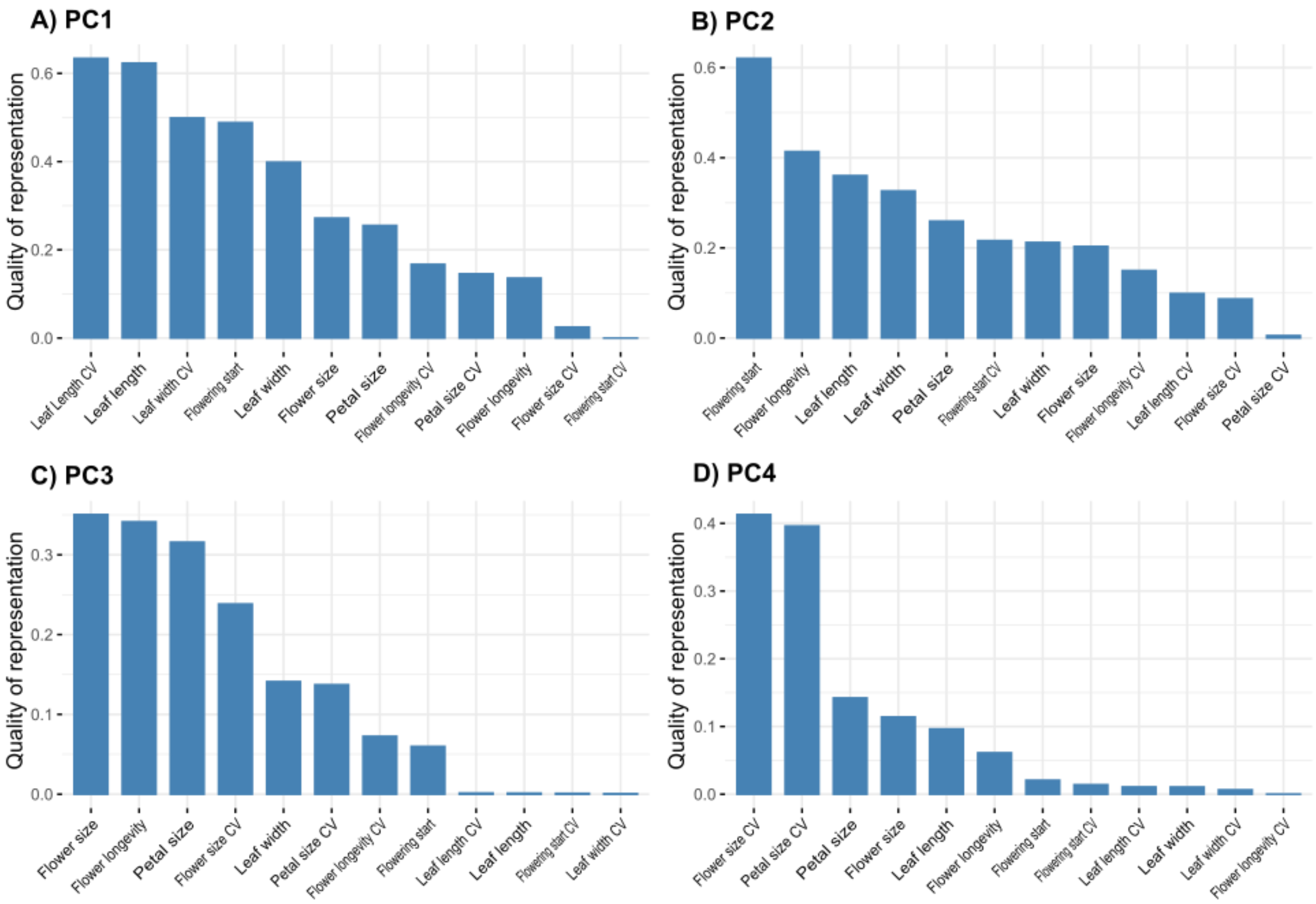


**Figure S1.** Contributions of the trait averages and trait coefficient of variation (CV) for A) principal component 1 (PC1); B) principal component 2 (PC2); C) principal component 3 (PC3); and D) principal component 4 (PC4) for the control treatment.

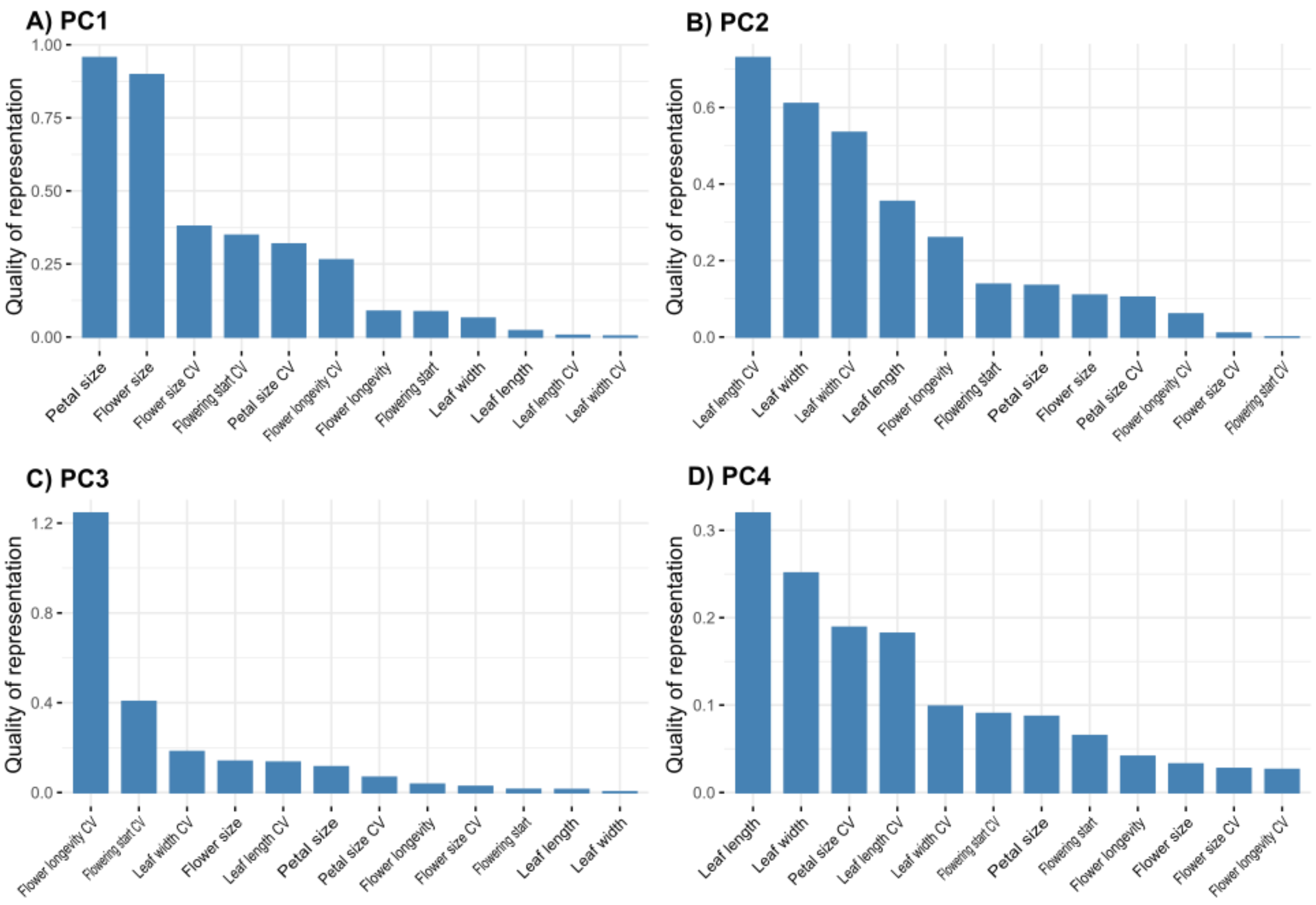


**Figure S2.** Contributions of the trait averages and trait coefficient of variation (CV) for A) principal component 1 (PC1); B) principal component 2 (PC2); C) principal component 3 (PC3); and D) principal component 4 (PC4) for the drought treatment.

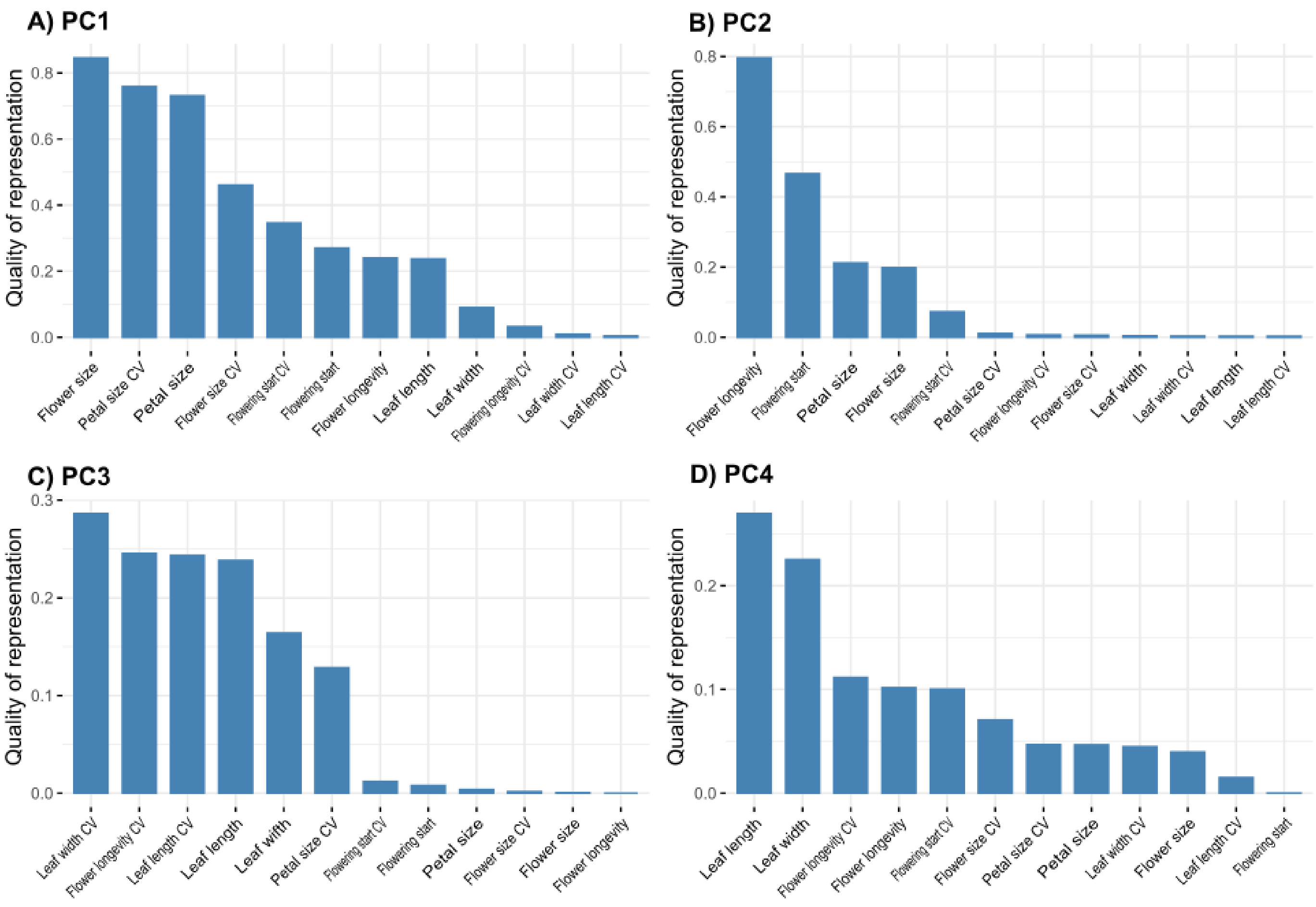


**Figure S3.** Contributions of the trait averages and trait coefficient of variation (CV) for A) principal component 1 (PC1); B) principal component 2 (PC2); C) principal component 3 (PC3); and D) principal component 4 (PC4) for the early shading treatment.

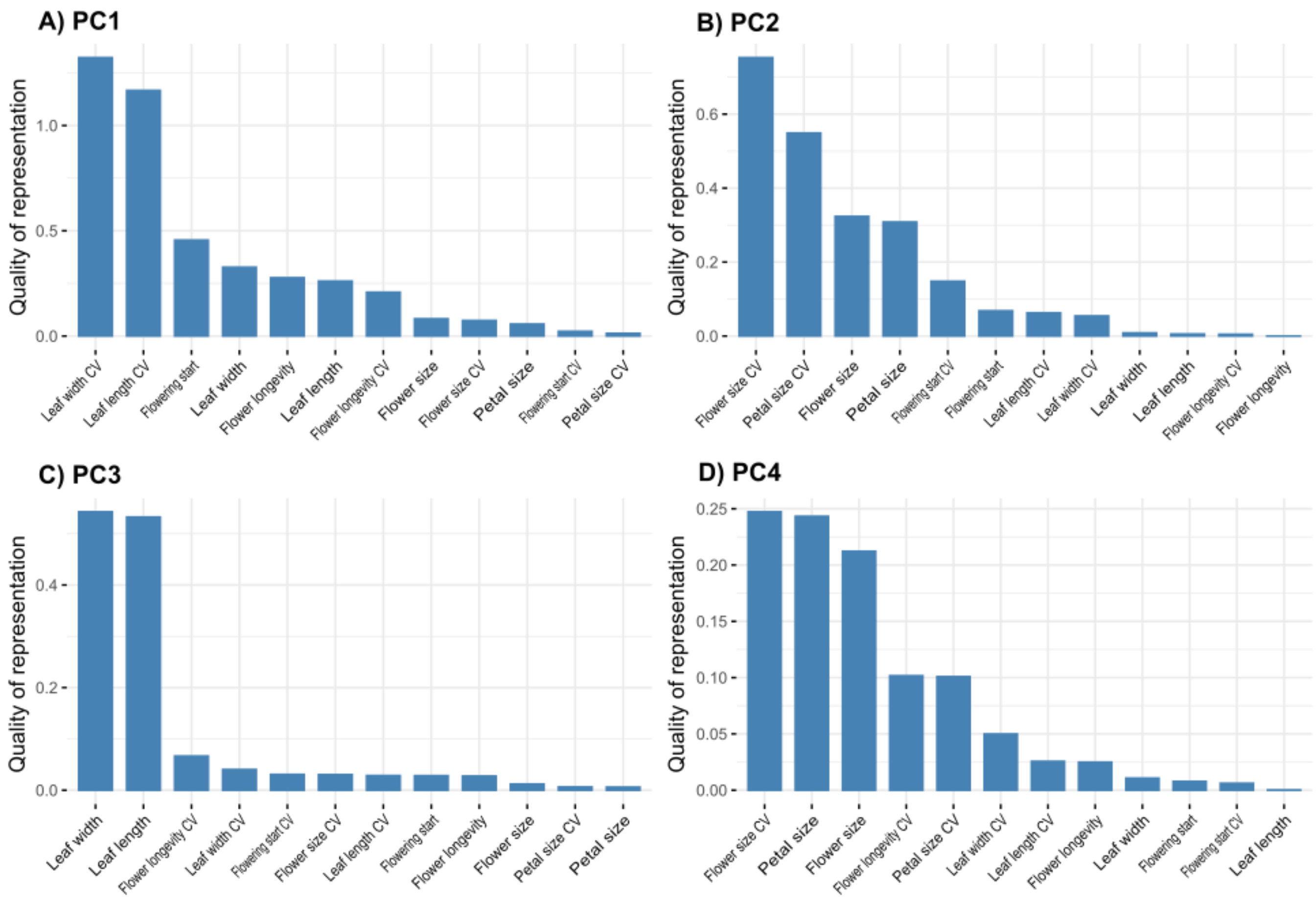


**Figure S4.** Contributions of the trait averages and trait coefficient of variation (CV) for A) principal component 1 (PC1); B) principal component 2 (PC2); C) principal component 3 (PC3); and D) principal component 4 (PC4) for the combined drought and early shading treatment.

**Table S1.** Overview of the distribution of the individuals included in the study design for the vegetative measurements (leaf length and leaf width), the treatments they were subjected to, the number of genets they originated from and the number of populations that had successfully been sampled in each of the three regions: Schwäbian Alb, Hainich-Dün, and Schorfheide-Chorin.

| | Populations (N) | Genets (N) | Treatments | Ramets |
|---|---|---|---|---|
| **Schwäbian Alb (ALB)** | 4 | 10 | Control | 14 |
| | | | Early Shading | 15 |
| | | | Drought | 35 |
| | | | Early Shading + Drought | 4 |
| **Hanich-Dün (HAI)** | 6 | 24 | Control | 48 |
| | | | Early Shading | 40 |
| | | | Drought | 50 |
| | | | Early Shading + Drought | 70 |
| **Schorfheide-Chorin (SCH)** | 5 | 17 | Control | 25 |
| | | | Early Shading | 60 |
| | | | Drought | 44 |
| | | | Early Shading + Drought | 20 |

| | | | | |
|---|---|---|---|---|
| **Total** | 15 | 51 | Control | 87 |
| | | | Early Shading | 115 |
| | | | Drought | 129 |
| | | | Early Shading + Drought | 94 |
| | | | **Total** | **425** |

**Table S2.** Overview of the distribution of the individuals included in the study design for the floral measurements (flowering start, flower longevity, flower size, and petal size), the treatments they were subjected to, number of genets they originated from and number of populations that had successfully been sampled in each of the three regions: Schwäbian Alb, Hainich-Dün, and Schorfheide-Chorin.

| | Populations (N) | Genets (N) | Treatments | Ramets |
|---|---|---|---|---|
| **Schwäbian Alb (ALB)** | 8 | 21 | Control | 168 |
| | | | Early Shading | 93 |
| | | | Drought | 202 |
| | | | Early Shading + Drought | 75 |
| **Hanich-Dün (HAI)** | 8 | 26 | Control | 147 |
| | | | Early Shading | 113 |
| | | | Drought | 132 |
| | | | Early Shading + Drought | 234 |
| **Schorfheide-Chorin (SCH)** | 5 | 17 | Control | 74 |
| | | | Early Shading | 183 |
| | | | Drought | 134 |
| | | | Early Shading + Drought | 66 |

| | | | | |
|---|---|---|---|---|
| **Total** | 21 | 64 | Control | 389 |
| | | | Early Shading | 389 |
| | | | Drought | 468 |
| | | | Early Shading + Drought | 375 |
| | | | **Total** | **1621** |

**Table S3. Effects of the applied treatments on trait-specific averages at the intra-ramet level.** Trait averages in *Galium odoratum* at the intra-ramet level calculated for each trait was used as response variables. Trait CV as well as the experimental treatments and their interaction were included as fixed factors and covariates. Genet nested within population and region was included as random factor. Significance levels are represented by asterisks: • *P* <0.1; * *P* < 0.05; ** *P* < 0.01; *** *P* < 0.001.

| Response | Predictor | Chisq | Df | P |
|---|---|---|---|---|
| Leaf length average | Trait CV | 11.729 | 1 | **<0.001 *** ** |
| | Shade | 3.510 | 1 | **0.061 •** |
| | Drought | 11.708 | 1 | **<0.001 *** ** |
| | Shade:Drought | 0.613 | 1 | 0.434 |
| Leaf width average | Trait CV | 14.340 | 1 | **<0.001 *** ** |
| | Shade | 3.890 | 1 | **0.049 * ** |
| | Drought | 9.659 | 1 | **0.002 ** ** |
| | Shade:Drought | 0.043 | 1 | 0.836 |
| Flowering start average | Trait CV | 0.897 | 1 | 0.345 |
| | Shade | 6.073 | 1 | **0.014 * ** |
| | Drought | 1.128 | 1 | 0.288 |
| | Shade:Drought | 1.246 | 1 | 0.264 |
| Flower longevity average | Trait CV | 22.541 | 1 | **<0.001 *** ** |
| | Shade | 1.200 | 1 | 0.273 |
| | Drought | 0.446 | 1 | 0.504 |
| | Shade:Drought | 0.280 | 1 | 0.597 |
| Flower size average | Trait CV | 9.647 | 1 | **0.002 ** ** |
| | Shade | 0.088 | 1 | 0.767 |
| | Drought | 1.046 | 1 | 0.306 |
| | Shade:Drought | 0.257 | 1 | 0.612 |
| Petal size average | Trait CV | 7.386 | 1 | **0.007 ** ** |
| | Shade | 0.082 | 1 | 0.775 |
| | Drought | 1.028 | 1 | 0.311 |
| | Shade:Drought | 0.712 | 1 | 0.399 |
